# Differential regulation of α-amino-3-hydroxy-5-methyl-4-isoxazolepropionic Acid (AMPA) receptor tetramerization by auxiliary subunits

**DOI:** 10.1101/2023.02.07.527516

**Authors:** Noele Certain, Quan Gan, Joseph Bennett, Helen Hsieh, Lonnie P. Wollmuth

## Abstract

AMPA receptor (AMPAR) auxiliary subunits are specialized, non-transient binding partners of AMPARs that modulate their ion channel gating properties and pharmacology, as well as their biogenesis and trafficking. The most well characterized families of auxiliary subunits are transmembrane AMPAR regulatory proteins (TARPs) and cornichon homologs (CNIHs) and the more recently discovered GSG1-L. These auxiliary subunits can promote or reduce surface expression of AMPARs in neurons, thereby impacting their functional role in membrane signaling. Here, we show that CNIH-2 enhances the tetramerization of wild type and mutant AMPARs, possibly by increasing the overall stability of the tetrameric complex, an effect that is mainly mediated by interactions with the transmembrane domain of the receptor. We also find CNIH-2 and CNIH-3 show receptor subunit-specific actions in this regard with CNIH-2 enhancing both GluA1 and GluA2 tetramerization whereas CNIH-3 only weakly enhances GluA1 tetramerization. These results are consistent with the proposed role of CNIHs as endoplasmic reticulum cargo transporters for AMPARs. In contrast, TARP γ-2, TARP γ-8, and GSG1-L have no or negligible effect on AMPAR tetramerization. On the other hand, TARP γ-2 can enhance receptor tetramerization but only when directly fused with the receptor at a maximal stoichiometry. Notably, surface expression of functional AMPARs was enhanced by CNIH-2 to a greater extent than TARP γ-2 suggesting that this distinction aids in maturation and membrane expression. These experiments define a functional distinction between CNIHs and other auxiliary subunits in the regulation of AMPAR biogenesis.

## Introduction

The majority of fast excitatory neurotransmission in the nervous system relies on AMPA receptors (AMPARs) (Groc & Choquet 2020, Hansen et al 2021). AMPARs are glutamate-gated ion channels that are regulated by the composition of the AMPAR macromolecular complex. The core of the complex is formed by pore-forming AMPAR subunits (GluA1-4). Functional properties of the core ion channel vary in a subunit-specific manner as well as by various post-translational modifications (e.g., RNA editing, glycosylation) (Shi et al 2001, Greger et al 2003, Lu & Roche 2012, Kandel et al 2018). In addition, this core AMPAR complex is typically associated with membrane-spanning auxiliary subunits (Jackson & Nicoll 2011, Yan & Tomita 2012, Jacobi & von Engelhardt 2020, Kamalova & Nakagawa 2020). Although these auxiliary subunits are not directly involved in the formation of the ion channel, they influence synaptic transmission by modulating AMPAR gating kinetics, ion permeation and channel block as well as pharmacological properties (Greger & Esteban 2007, Jackson & Nicoll 2011, Haering et al 2014, Bissen et al 2019). AMPAR auxiliary subunits also regulate surface expression and distribution of receptors (Sumioka 2013, Khodosevich et al 2014, Bissen et al 2019). Auxiliary subunits associate with AMPARs in a neuronal and subunit specific-dependent manner, which contributes greatly to the functional diversity of AMPARs in the mammalian central nervous system (Shi et al 2009, Schwenk et al 2014, McGee et al 2015, Jacobi & von Engelhardt 2017).

TARPs and CNIHs are the most abundant AMPAR auxiliary subunits in the brain (Schwenk et al 2014). Similar to TARPs, cornichon-2 (CNIH-2) and cornichon-3 (CNIH-3) enhance AMPAR channel conductance and slow receptor deactivation and desensitization (Schwenk et al 2009, Shi et al 2010, Coombs et al 2012) despite a distinctive evolutionary origin. CNIH-2 also shows enrichment in both the ER and synaptic compartments (Schwenk et al 2019).

CNIHs function as endoplasmic reticulum (ER) cargo exporters, mediating AMPAR maturation and translocation from the ER and golgi complex to the surface membrane (Schwenk et al 2009, Harmel et al 2012, Brockie et al 2013). Neurons lacking CNIH-2/-3 show severe reduced synaptic AMPARs activity (Herring et al 2013). Knockdown of CNIH-2 results in the retention of AMPARs and therefore disruption of synaptic functions (Herring et al 2013).

Stargazin (TARP γ-2), the prototypical member of the TARPs family, is the most extensively studied AMPAR auxiliary subunit. TARP γ-2 increases AMPAR agonist potency and efficacy, channel conductance, and slows AMPAR deactivation and desensitization (Priel et al 2005, Cho et al 2007, Milstein et al 2007). In addition to the impact of TARP γ-2 on receptor functions, TARP γ-2 also increases AMPAR surface expression and participates in synaptic targeting and receptor endocytosis, which underlies its implication in various forms of synaptic plasticity (Bedoukian et al 2006, Matsuda et al 2013, Constals et al 2015). Under certain conditions such as ER stress, TARP γ-2 promotes the delivery of AMPARs to the surface, but in a less efficient manner compared to CNIH-2 (Vandenberghe et al 2005, Shanks et al 2010, Schwenk et al 2019). TARP γ-2 plays an essential role in receptor surface trafficking (Chen et al 2000, Kato et al 2010b), but may come into play at later stages of receptor maturation (Shanks et al 2010).

TARP γ-8, another prominent TARP preferentially expressed in the hippocampus, which regulates AMPAR trafficking and synaptic density (Rouach et al 2005, Sumioka et al 2011). The knockdown of TARP γ-8 in primary hippocampal neurons leads to gross reduction in surface AMPAR expression (Zheng et al 2015). GSG1-L is a more recently identified AMPAR auxiliary subunit, which is structurally similar to the TARP family (Shanks et al 2012, Twomey et al 2017). GSG1-L reduces AMPAR surface expression by facilitating the endocytosis of surface AMPARs (Gu et al 2016, Kamalova et al 2020). Although auxiliary subunits are involved in receptor trafficking, mechanisms involved in these processes are incompletely understood.

In the present study, we explored the role of auxiliary subunits in regulating AMPAR tetramerization, an early step in receptor maturation (Greger et al 2003, Salussolia et al 2013, Morise et al 2019). We find that CNIH-2 and CNIH-3 enhance wildtype AMPAR tetramerization by increasing the number of available tetramers. This effect is likely mediated by interactions between CNIHs and the transmembrane domain (TMD) of AMPARs. In contrast, other prominent auxiliary subunits including TARP γ-2 and γ-8 and GSG1-L have no notable effect on AMPAR tetramerization, possibly due to late-stage association with AMPARs in the ER. We therefore conclude CNIHs differ in their ability to modulate AMPAR tetramerization based on the stage of involvement with AMPAR biogenesis in comparison to other AMPAR auxiliary subunits.

## Materials and Methods

### Molecular biology and heterologous expression

AMPA receptor (AMPAR) cDNA constructs are rat (*Rattus norvegicus*) flip splice variants for GluA1 (#P19490) and the edited (R) form of GluA2 (#P19491). Mutations were introduced in these constructs using PCR-based site-directed mutagenesis methods and were validated by DNA sequencing. Numbering is for the mature AMPAR protein with signal peptides: GluA1, 18 residues; GluA2, 21 residues. The GluA1-ΔCTD, HA-GluA1-ETD-EGFP, and GluA1-EGFP (C-terminal) were generated as described previously(Salussolia et al 2013, Gan et al 2016). The cDNA constructs were provided by: Dr. David Bredt (Johnson & Johnson), TARP γ-2-EYFP; Dr. Bernd Fakler (Freiburg), human CNIH-2 (Q6PI25) and GSG1-L (Q6UXU4); Dr. Stefan Herlitze (Ruhr), rat TARP γ-2 (Q71RJ2); Dr. Roger Nicoll (UCSF), GluA1-TARP γ-2 tandem; and Susumu Tomita (Yale), human CNIH-3 (Q8TBE1) and mouse TARP γ-8 (Q8VHW2). The GluA1 tandem contains GluA1 co-joined at the C-terminus with a short linker sequence to the N-terminus of rat TARP γ-2 (Shi et al 2009).

For BN-PAGE and whole cells recordings, human embryonic kidney (HEK) 293 cells (ATCC, CRL-1573) and HEK 293T (ATCC, CRL-3216) were transiently transfected with cDNAs using Xtreme Gene HP (Roche). For ICC, Neuro-2a (N2a) (ATCC, CCL-131) were transiently transfected with cDNAs using Xtreme Gene HP (Roche). AMPARs and auxiliary subunits were co-transfected at a cDNA ratio of 1:1. Transfected cells were maintained in media containing 5% FBS supplemented with 10 μM CNQX (Sigma) and 10 μM NBQX (Sigma) to minimize AMPAR-mediated excitotoxicity (Menuz et al 2007).

### Blue native-PAGE

HEK 293 cells were plated on 60 mm tissue culture dishes and transfected 24 hours later at 70% confluency. From 30-48 hours after transfection, the cells were harvested as previously described (Salussolia et al 2016). Membrane proteins extracted from whole-cell lysates were resolved using Blue Native-PAGE (BN-PAGE) as previously described (Schagger et al 1994, Gan et al 2016, Salussolia et al 2016). Briefly, protein samples mixed with 1X Native PAGE sample buffer, 0.05% NativePAGE G-250 additive, and 1% n-Dodecyl-β-D-maltoside were loaded onto Novex 4-16% Bis/Tris gradient gels (Life Technologies). When assaying stoichiometry of AMPAR-auxiliary subunit complexes, 3-12% Bis-tris gels were used in order to better resolve protein complexes at higher molecular masses. Molecular mass markers were also loaded and included NativeMARK Unstained Protein Standard (Life Technologies) and Apoferritin (Sigma). Blue Native-PAGE and transfer were conducted as previously described(Gan et al 2016, Salussolia et al 2016). BN-PAGE samples were analyzed by western blot. Anti-GluA1 (Millipore, MAB2269, 1:1500) and Anti-GluA2 (Millipore; MAB297, 1:500) mouse monoclonal antibodies were used to detect specific AMPAR subunits. HRP-conjugated anti-mouse IgG (Santa Cruz Biotechnologies, sc-2031, 1:1000) was used as a secondary antibody. Blots were developed using luminol reagent (Santa Cruz Biotechnologies, sc-2048) before exposure to chemiluminescence blue-sensitive film. For clarity of presentation, reorganized lanes from the same gel are indicated by a thin space between lanes.

### BN-PAGE densitometry and quantification

Developed immunoblot films were scanned using an EPSON flat scanner (Epson Perfection V700 Photo) into .tiff format in an 8-bit gray scale mode, 300 dpi resolution, and quantified using NIH Image J (Schneider et al 2012). The following parameters including antibody concentration, duration of immunostaining, and time of exposure were optimized to ensure major bands fall within the linear response range of the film (Salussolia et al 2016). Quantification of each resolved band for any oligomeric state (monomer, dimer, and tetramer bands) of AMPARs is based on the relative position to the molecular weight markers. Using ImageJ, the relative measurement of fraction of tetramer intensity is quantified as previously described (Gan et al 2016, Salussolia et al 2016). The tetramer fraction data was normalized to 1. The intensity of each band is not necessarily proportional to the total amount of protein within each band, because primary antibodies are not guaranteed to bind every single AMPAR subunit in an oligomeric complex. The fraction of tetramer quantification is therefore a relative measurement of tetramerization efficiency and cannot calculate absolute physical quantities as previously shown (Schagger et al 1994). Comparisons were made within the same cell passage due to variation in gene expression depending on the passage number.

### Whole-cell current recordings

HEK 293T cells were plated in 24-well plates at a density of 4 × 10^5^ cells on uncoated coverslips. Cells were maintained in 10% FBS at 37°C and 95% O_2_/5% CO_2_. AMPAR-mediated currents in transfected HEK 293T cells were recorded 24 - 48 hours following transfection. On each recording day, we recorded cells transfected with GluA1, GluA1 + CNIH-2, and GluA1 + TARP γ-2. Recordings were performed in the whole-cell configuration at room temperature (20-23°C) using an EPC-10 amplifier and Patchmaster software (HEKA Elektronik, Lambrecht, Germany). Signals were low-pass filtered at 5 kHz and digitized at 20 kHz. Patch microelectrodes were filled with a KCl-based intracellular solution (in mM): 140 KCl, 2 NaCl, 4 Mg^2+^-ATP, 0.3 Na^+^ -ATP, 1 BAPTA, 10 HEPES, pH 7.2 (KOH), 300 mOsm (sucrose) and had resistances of 4-6 MΩ. We did not use series resistance compensation, nor did we correct for junction potentials. External solution consisted of the following (in mM): 140 NaCl, 1.8 CaCl_2_, 1 MgCl_2_, and 10 HEPES, pH = 7.2 (NaOH). All solutions contained 15 mM of cyclothiazide (CTZ) to minimize the impact of desensitization on current amplitudes. Glutamate was applied using a piezo-driven double barrel application system with one barrel containing external solution and the other containing the same solution with added glutamate (3 mM) (Yelshansky et al 2004, Amin et al 2018). The 10-90% rise time of the application pipet was approximately 500 - 800 μs.

We measured membrane capacitance (*C*_*m*_), maximum current amplitudes (*I*_*peak*_), and the current-voltage relationship. The current amplitude is a function of the number of ion channels on the membrane (*N*), the probability of opening (*P*_*open*_), and their single channel conductance (γ):

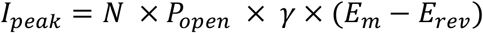

Where membrane potential (*E*_*m*_) and reversal potential (*E*_*rev*_) are assumed to be invariant in our experiment. CNIH-2 and TARP γ-2 have differential effects on *P*_*open*_ and γ (Coombs et al 2012). Since we were interested in comparing the effect of CNIH-2 and TARP γ-2 on the number of channels on the membrane, we calculated *N* using our current amplitudes and previously published values for *P*_*open*_ and γ using the relationship:

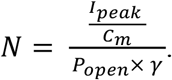

Values for *P*_*open*_ and γ were: 0.46 and 30.2 pS (GluA1 + CNIH-2) and 0.64 and 28.2 pS (GluA1 + TARP γ-2), while the values for wildtype GluA1 alone were 0.41 and 18.7 pS (Coombs et al 2012). Note we do not view the derived *N* as absolute, but rather as a relative index of channels.

### Immunocytochemistry

N2a cells were transfected with GluA1(G816A)-eGFP (C-terminal) and auxiliary subunits at a 1:1 DNA ratio. Post 48-hour transfection, transfected N2a cells were stained with anti-GluA1 antibody (MilliporeSigma #MAB2263; 1:500) in conditioned media for 30 minutes at 37°C. Coverslips were washed with 1X phosphate buffered saline (PBS), and then fixed for 10 minutes with 4% paraformaldehyde and 4% sucrose in 1X PBS, pH 7.4. The coverslips were blocked with blocking solution (5% normal goat serum in 1X PBS) for 1 hour at 4°C. Coverslips were washed with PBS and then incubated with Alexa 633-conjugated goat anti-mouse secondary antibody (Invitrogen #A21052; 1:1500) for 1 hour at 4°C. After washing with PBS, coverslips were stained with NucBlue Ready Probes (Invitrogen #R37606) for 20 minutes and mounted in Prolong Diamond mount (Invitrogen #P36961).

Images were acquired on an Olympus FV-1000 confocal microscope with an oil-immersion objective (60x, numerical aperture 1.42) using a Fluoview software (FV10-ASW, version 4.02, Olympus). For each coverslip, 3-5 fields were imaged with consistent filter settings. For quantification of surface AMPARs, corrected total cell fluorescence (CTCF) was determined on ImageJ (NIH) by measuring the surface AMPARs (Alexa 633) or total AMPARs (GFP tag) integrated density for the N2a cell area and subtracting off-cell background mean intensity. Intensity was then normalized to the mean intensity of GluA1(G816A) when expressed alone. Quantification, imaging, and image analysis were done blind to treatment conditions. Images collected were blinded using Blind Analysis Tools, an ImageJ plugin.

### Statistics

Data analysis for BN-PAGE, whole-cell recordings, and immunocytochemistry were performed using Igor Pro (version 7, Wave Metrics) and Microsoft Excel Analysis ToolPak. All data values are represented as the mean ± SEM. To ensure reproducibility of results at least three independent experiments were performed. Analyses of tetramer fraction, current amplitudes, relative channel index, and fluorescence intensity were performed between different constructs or combinations of co-expression using *t-test* or a *one-way ANOVA post hoc Tukey’s test*.

## Results

To address the role of AMPAR auxiliary subunits in receptor tetramerization, we expressed AMPARs either alone or in the presence of different auxiliary subunits. We used BN-PAGE to evaluate the oligomeric states of AMPARs in whole-cell lysates. Initially, we assayed GluA1 tetramerization.

### CNIHs uniquely promote GluA1 tetramerization

GluA1 expressed alone readily oligomerizes, forming predominantly tetramers with a small albeit notable dimer band and generally no monomer bands (**Figures 1A-E, *left panels***) (Greger et al 2003, Penn et al 2008, Rossmann et al 2011, Salussolia et al 2013, Gan et al 2016). In the presence of CNIH-2, GluA1 tetramerization was nearly complete, with negligible amount of dimer remaining (**Figure 1A, *left panel***). To quantify these results, we derived the tetramer fraction from cumulative oligomeric band intensities (see Materials & Methods). Although GluA1 alone showed a high tetramer fraction, when co-expressed with CNIH-2 the protein fraction was now almost completely tetrameric (**Figure 1A, *right panel***).

**Figure 1.**
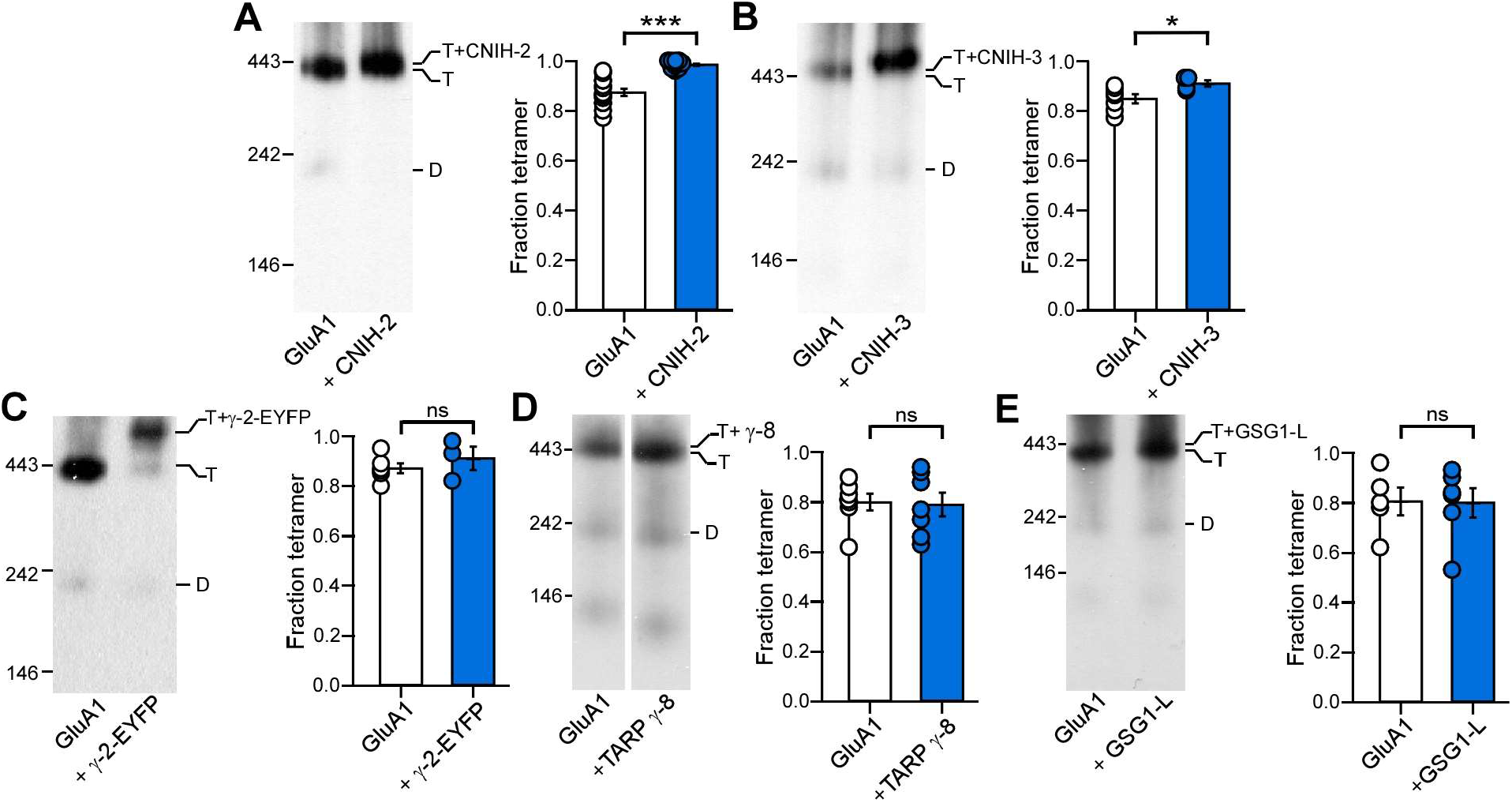
Cornichon homologs promote assembly of GluA1 tetramers. (**A-E**) BN-PAGE (*left panels*) and normalized tetramer fractions (*right panels*) of GluA1 expressed alone or with auxiliary subunits CNIH-2 (**A**), CNIH-3 (**B**), TARP γ-2-EYFP (γ-2-EYFP) (**C**), TARP γ-8 (**D**), or GSG1-L (**E**). Normalized tetramers fraction in this figure and all subsequent figures are shown as mean ± SEM. Constructs were expressed in HEK 293 cells. For BN-PAGE gels, positions of molecular mass markers are labeled on the left (see Methods). Oligomeric states of the detected bands were estimated by molecular mass and indicated as T (tetramers) or D (Dimers). For bar plots, circles represent individual values. Number of gels (left to right) and significance (*t-test*) of tetramer fraction relative to control: 13, 10, *p <0*.*001* (**A**); 7, 4, *p = 0*.*03* (**B**); 6, 3, *p = 0*.*52* (**C**); 7, 7, *p = 0*.*84* (**D**); 5, 6, *p = 0*.*95* (**E**). In plots, significance is indicated (*\*p < 0*.*05* or *\*\*\*p < 0*.*001; ns*, not significant).

Within the cornichon homologs, CNIH-2 shares a unique sequence with CNIH-3, which carries specificity for binding to AMPARs (Schwenk et al 2009, Shanks et al 2012, Shanks et al 2014). When co-expressed with homomeric GluA1 (**Figure 1B**), CNIH-3 led to a significant increase in the tetramer fraction, like CNIH-2, although a small dimer population remained. In contrast to CNIHs, other AMPAR auxiliary subunits including TARP γ-2 (**Figure 1C**), TARP γ-8 (**Figure 1D**) and GSG1-L (**Figure 1E**) showed no significant effect on receptor tetramerization. Overall, these results suggest that CNIHs have the unique function to enhance AMPAR tetramerization.

### CNIHs can rescue disrupted AMPAR tetramerization

Interactions between the M4 transmembrane helices and the ion channel core (M1-M3) are required to assemble a functional AMPAR (Salussolia et al 2011, Salussolia et al 2013, Gan et al 2016, Amin et al 2017). Within the M4 segment are key residues, the ‘VVLGAVE’ motif, that directly interact with the ion channel core (Sobolevsky et al 2009). Amino acid substitutions at these positions leads to altered receptor tetramerization (Salussolia et al 2013, Amin et al 2017).

To further test whether CNIHs promote the tetramerization process in comparison to other AMPAR auxiliary subunits, we co-expressed auxiliary subunits with GluA1 AMPARs containing a single-point mutation, a glycine (G) to alanine (A), in the ‘VVLGAVE’ motif. Despite the subtle difference in the side chain, G816A in GluA1 attenuates tetramer formation (**Figure 2, *left panel***) to about 50% assayed using either BN-PAGE (**Figure 2A, *right panel***) or fluorescent size exclusion chromatography (FSEC) (Amin et al 2017). Notably, co-expression with CNIH-2 fully rescued the tetramerization deficit of GluA1(G816A) to wild type levels (**Figure 2A**). CNIH-3 also rescued the GluA1(G816A) tetramerization deficit, although to a lesser extent (**Figure 2B**). In contrast, and as observed for wild type GluA1, co-expression of TARP γ-2 (**Figure 2C**), TARP γ-8 (**Figure 2D**), or GSG1-L (**Figure 2E**) with GluA1(G816A) had no significant effect on the tetramer fraction. In summary, CNIHs but most notably CNIH-2 has a strong effect on the assembly process of GluA1, whereas other AMPAR auxiliary subunits do not.

**Figure 2.**
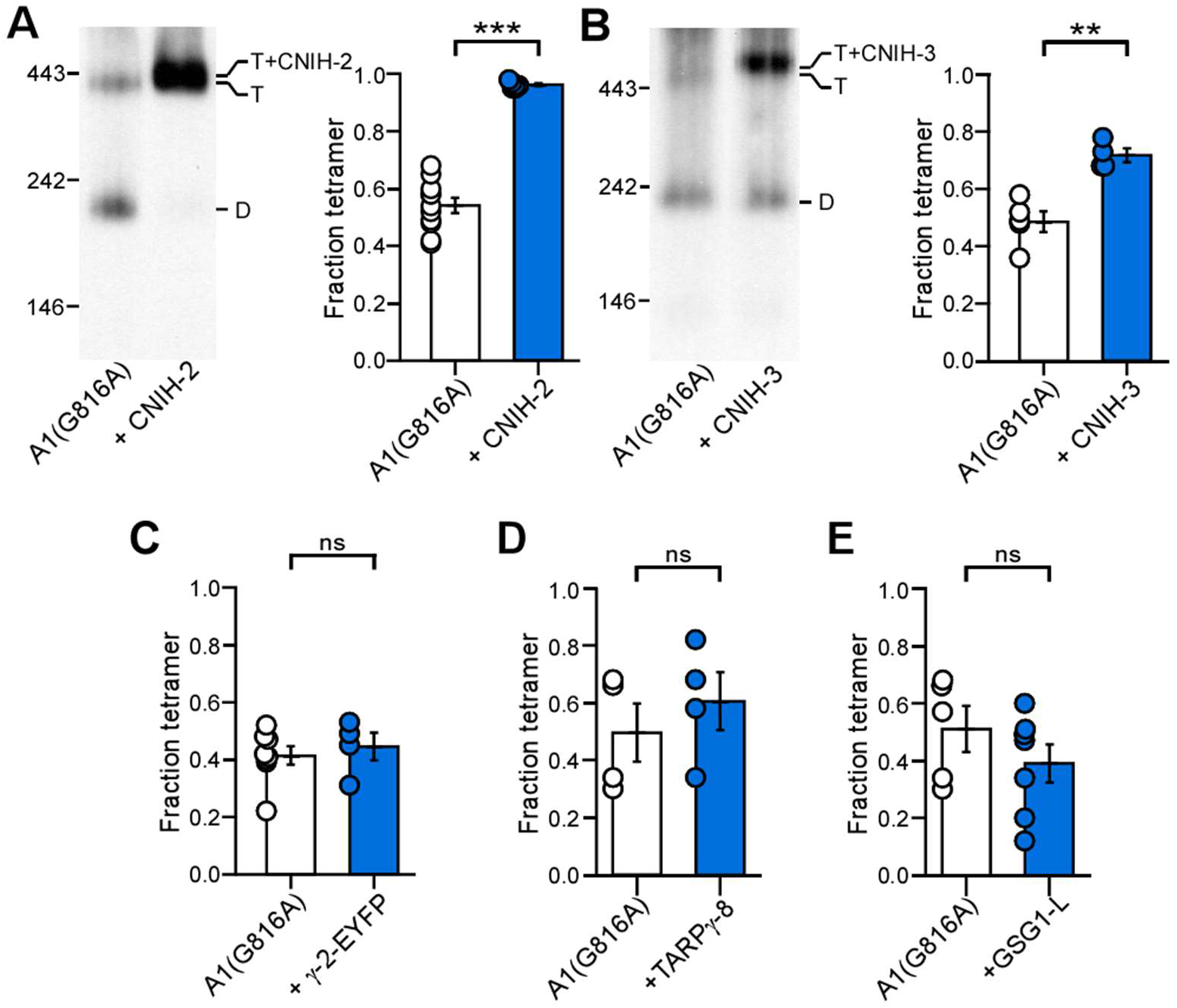
Cornichon homologs can rescue deficits in AMPAR tetramerization. (**A-B**) BN-PAGE (*left panels*) and normalized tetramer fractions (*right panels*) of GluA1(G816A) expressed alone or with CNIH-2 (**A**) or CNIH-3 (**B**). G816A is a mutation in the M4 VVLGAVE face that attenuates receptor tetramerization. (**C-E**) Normalized tetramer fractions of GluA1(G816A) expressed alone or with TARP γ-2-EYFP (γ-2-EYFP) (**C**), TARP γ-8 (**D**), or GSG1-L (**E**). Number of gels (left to right) and significance (*t-test*) of tetramer fraction relative to control: 11, 6, *p <0*.*001* (**A**); 5, 4, *p = 0*.*0014* (**B**); 8, 4, *p = 0*.*61* (**C**); 4, 4, *p = 0*.*46* (**D**); 5, 7, *p = 0*.*28* (**E**). In plots, significance is indicated (*\*\*p < 0*.*01* or *\*\*\*p < 0*.*001; ns*, not significant).

### CNIH-2 distinctively enhances GluA2(R) tetramerization

Like GluA1, the GluA2 AMPAR subunit is prominently expressed throughout the brain and interacts with a variety of auxiliary subunits (Schwenk et al 2014, Zhao et al 2019). GluA2 exists exclusively in an edited form, indicated as GluA2(R), where an asparagine (N) is replaced with an arginine (R) in the mature protein (Hansen et al 2021). We therefore tested the effects of auxiliary subunits on GluA2(R) tetramerization.

Like GluA1, homomeric GluA2(R) efficiently forms tetramer (**Figures 3A, 3C, & 3E**) (Greger et al 2003, Pick & Ziff 2018). When co-expressed with CNIH-2, wild type GluA2(R) showed enhanced tetramerization similar to the effects on GluA1 (**Figure 3A**). In contrast, co-expression of CNIH-3 with GluA2(R) showed no change to the tetramer fraction (**Figure 3C**), suggesting a differential effect between CNIH-2 and CNIH-3 on GluA2(R) tetramerization. To further verify these differential effects, we assayed the G-to-A substitution (VVLGAVE) in GluA2(R) that attenuates tetramerization (**Figures 3B & 3D**). Consistent with what is observed for wild type GluA2(R), the tetramerization process was recovered by co-expression with CNIH- 2 (**Figure 3B**), but not with CNIH-3 (**Figure 3D**). Additionally, TARP γ-2 had no significant effect on GluA2(R) tetramerization, either for wild type GluA2(R) (**Figure 3E**) or for GluA2(R)(G823A) (**Figure 3F**).

**Figure 3.**
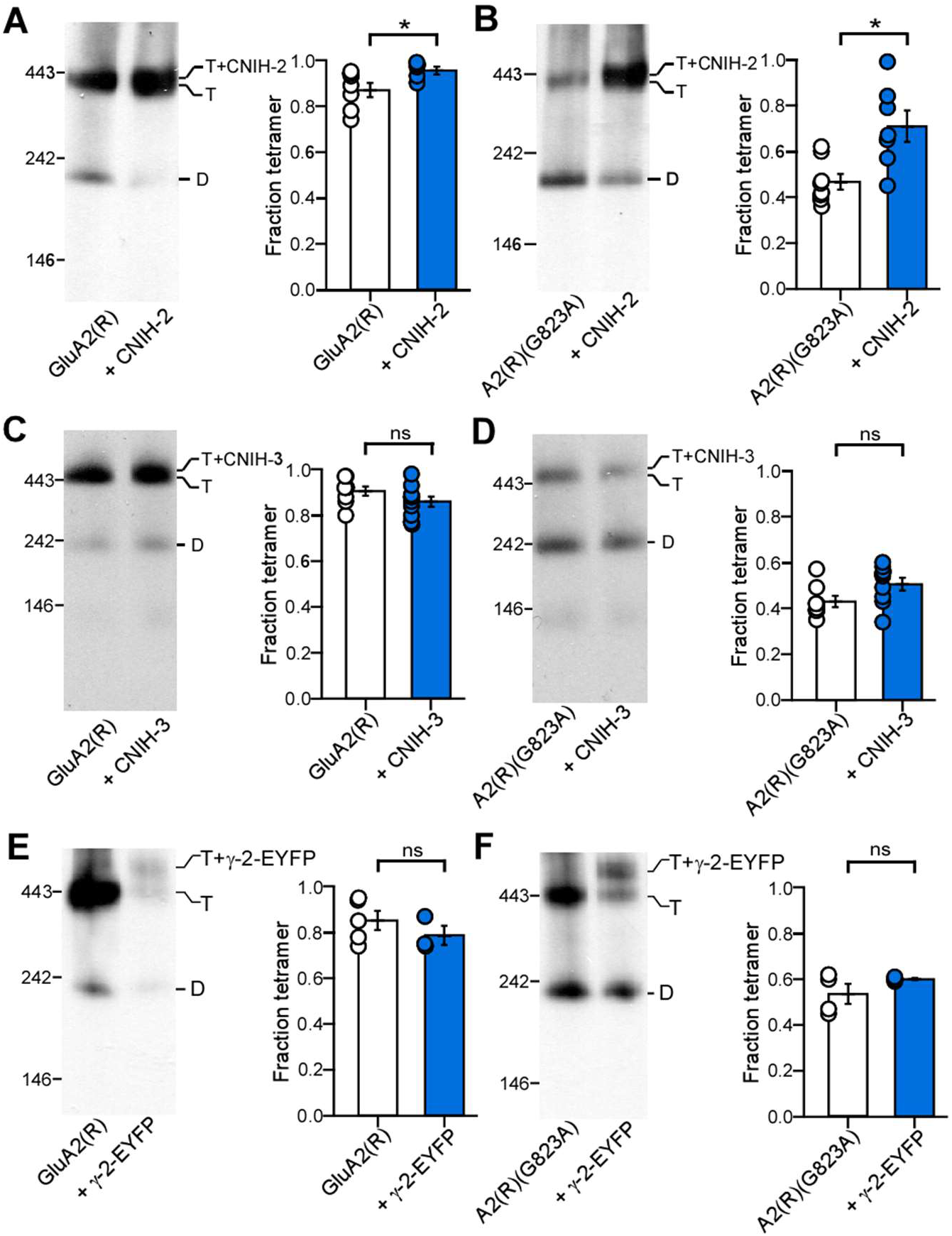
Cornichon homologs differentially regulate GluA2 assembly. (**A-F**) BN-PAGE (*left panels*) and normalized tetramer fractions (*right panels*) of GluA2(R) (**A, C, E**) or GluA2(R)(G823A) (**B, D, F**) expressed alone or with indicated auxiliary subunits. G823A is a mutation in the M4 VVLGAVE face that attenuates receptor tetramerization. Number of gels (left to right) and significance (*t-test*) of tetramer fraction relative to control: 7, 5, *p = 0*.*041* (**A**); 8, 7, *p = 0*.*011* (**B**); 8, 10, *p = 0*.*14* (**C**); 8, 9, *p = 0*.*068* (**D**); 5, 3, *p = 0*.*31* (**E**); 4, 4, *p = 0*.*22* (**E**) In plots, significance is indicated (*\*p < 0*.*05; ns*, not significant).

The results with the G-to-A substitution in the ‘VVLGAVE’ face strongly supports the differential role of CNIH-2 in enhancing GluA1 (**Figure 2**) and GluA2(R) (**Figure 3**) tetramerization. However, it is possible that a single substitution within the ‘VVLGAVE’ motif can specifically disrupt the function of TARP γ-2, but not that of CNIHs. We therefore tested other subtle mutations in the ‘VVLGAVE’ motif where the side chain replacement was similar to the endogenous residues: in GluA1(V823L) and (E827D); and in GluA2(R)(V830L) and (E834D). These substitutions in GluA1 attenuate tetramerization (Amin et al 2017). Based on BN-PAGE, these substitutions produce moderate tetramerization deficits that were rescued by CNIH-2 co-expression, but not by TARP γ-2 co-expression (**Figure 4**). Thus, the differential effects between CNIH-2 and other auxiliary subunits do not depend on a particular mutation within the VVLGAVE motif but applies to the AMPAR tetramerization process in general.

**Figure 4.**
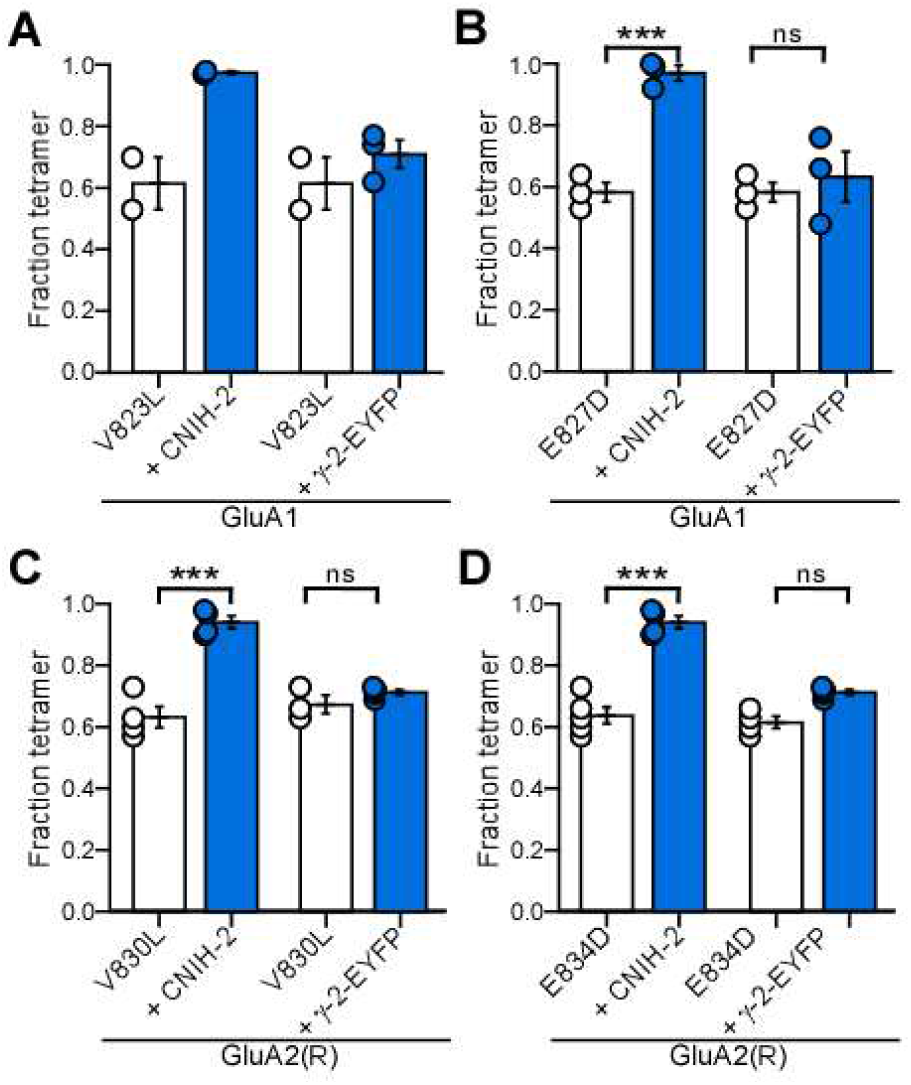
Rescue of AMPAR tetramerization by CNIH-2 is independent of the position altered in the VVLGAVE motif. Normalized tetramer fractions for single substitutions in the VVLGAVE motif in GluA1 (**A & B**) or GluA2(R) (**C & D**) expressed alone or with indicated auxiliary subunits. Constructs that disrupt tetramerization include: GluA1(V823L) (**A**), GluA1(E827D) (**B**), GluA2(R)(V830L) (**C**), and GluA2(R)(E834D) (**D**). Number of gels (left to right) and significance (*t-test*) of tetramer fraction relative to control: 2, 2, 2, 3, *p values not determined* (**A**); for CNIH-2: 3, 3, *p < 0*.*001* and for γ-2: 3, 3, *p = 0*.*62* (**B**); for CNIH-2: 4, 4, *p < 0*.*001* and for γ-2: 4, 3, *p = 0*.*16* (**C**); for CNIH-2: 5, 4, *p < 0*.*001* and for γ-2: 5, 4, *p = 0*.*06* (**D**). In plots, significance is indicated (*\*\*\*p < 0*.*001; ns*, not significant).

### CNIH-2 acts through the AMPAR transmembrane domain to promote assembly

Although the M4 segment substitutions disrupt tetramerization and CNIH-2 can rescue these deficits (**Figures 2-4**), these results do not indicate that CNIHs interacts with the M4 segment to facilitate tetramerization. AMPARs display a modular structure consisting of four major structural domains: extracellular amino terminal (ATD) and ligand-binding (LBD) domains, a transmembrane domain (TMD) forming the ion channel, and an intracellular carboxyl terminal domain (CTD) (**Figure 5A**). To assess which of these domains mediate the effect of CNIH-2 on AMPAR tetramerization, we used constructs with entire domains deleted from GluA1 (Gan et al 2016).

**Figure 5.**
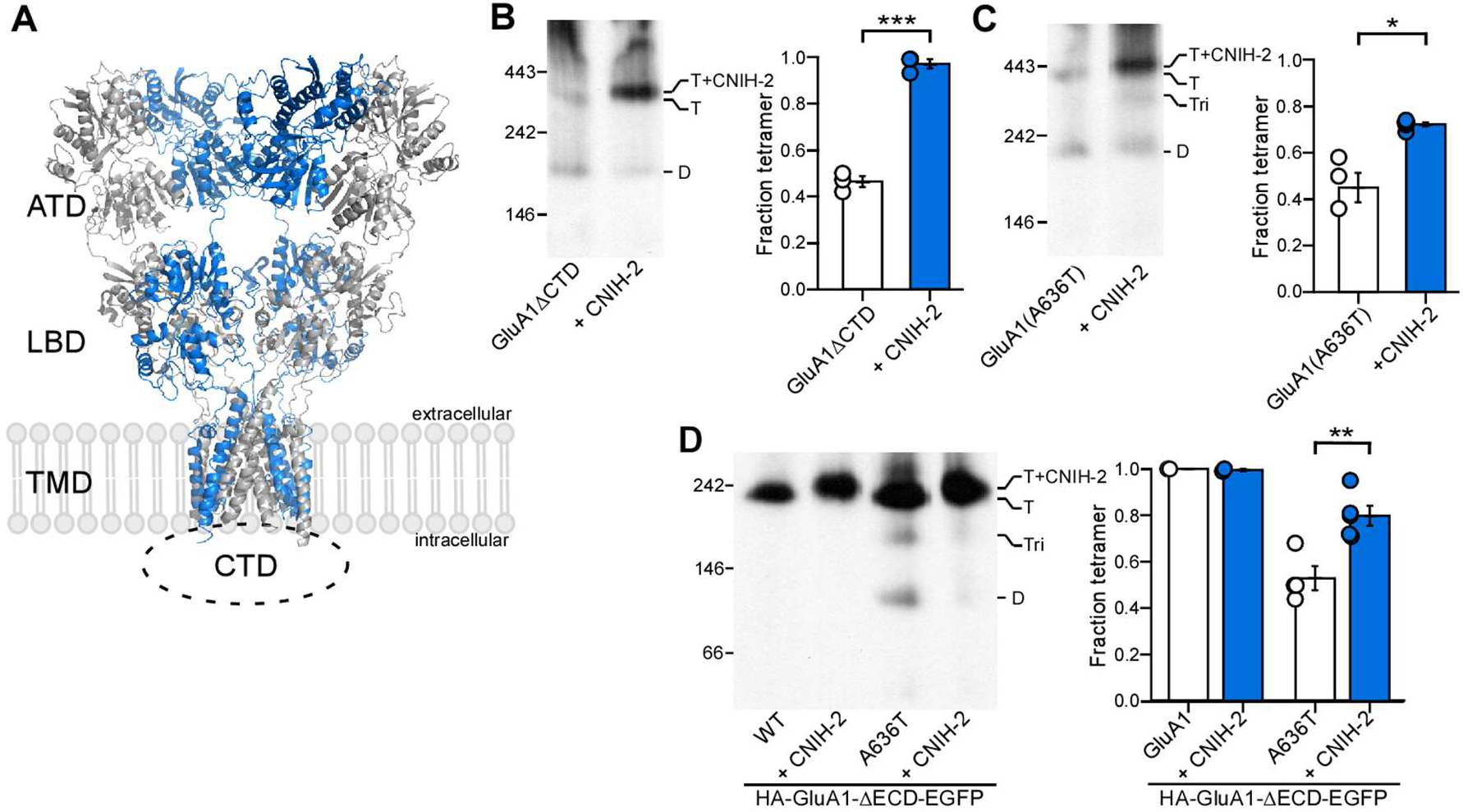
CNIH-2 acts through the transmembrane domain to promote tetramerization. (**A**) Structure of AMPARs consist of four modular domains: Extracellular amino-terminal (ATD) and ligand-binding (LBD) domains, the transmembrane domain (TMD) forming the ion channel, and the intracellular C-terminal domain (CTD). Homomeric GluA2 tetramer, subunit A’ C’ pairs in gray and B’D’ pairs in blue; PDB:6SKX (Yelshanskaya et al 2020) (**B-D**) BN-PAGE (*left panels*) and normalized tetramer fractions (*right panels*) of GluA1 lacking the C-terminal domain (ΔCTD) (**A**); GluA1 containing the lurcher mutation (A636T) (**B**); and GluA1 lacking the extracellular domains (ΔECD) (**C**). Number of gels (left to right) and significance (*t-test*) of tetramer fraction relative to control: 3, 3, *p < 0*.*001* (**B**); 4, 5, *p = 0*.*013* (**C**); for GluA1: 2, 2, *p value not determined* and for A636T: 4, 5, *p = 0*.*007* (**D**). In plots, significance is indicated (*\*p < 0*.*05, **p < 0*.*01*, or *\*\*\*p < 0*.*001; ns*, not significant).

The CTD of AMPAR is involved in receptor trafficking and regulation of receptor functions and AMPAR subunit-dependent (Tomita et al 2005, Granger et al 2013, Jenkins et al 2014, Zhou et al 2018, Diaz-Alonso et al 2020). The AMPAR CTD plays a critical role in physical interaction with auxiliary subunits and in enabling their modulatory effects on receptor function (Ben-Yaacov et al 2017). As found previously, complete deletion of GluA1 (GluA1-ΔCTD) showed a deficit in tetramerization, but this deficit is completely reversed by CNIH-2 co-expression (**Figure 5B**) (Gan et al 2016). Hence, the GluA1 CTD is not required for either the association of CNIH-2 with the receptor or its effect on tetramerization.

Testing the role of the other domains is non-trivial. However, we took an approach to address this question where the cumulative outcome indicates that CNIH-2 exerts its action via the TMD. Initially, we introduced the ‘lurcher’ mutation A636T into GluA1 and co-expressed with CNIH-2. The position, A636, is in the highly conserved ‘SYTANLAAF’ motif of the transmembrane domain (**Figure 5A**) and may disrupt tetramer assembly due to its involvement in the “dimerization of dimers” process (**Figure 5C)** (Kim et al 2010). As shown previously, the lurcher mutation in GluA1 reduces receptor tetramerization (Kim et al 2010, Gan et al 2016). Notably, the co-expression of CNIH-2 with the GluA1 lurcher mutant led to a significant increase in the tetramer fraction (**Figure 5C**), indicating that CNIH-2 can overcome the disruption of the lurcher mutation on tetramerization.

The function of auxiliary subunits has been tied to contacts with the ATD and/or the LBD (Cais et al 2014, Riva et al 2017, Klykov et al 2021, Watson et al 2021, Herguedas et al 2022). To evaluate whether the extracellular domains (ECDs), the ATD and the LBD, are required for CNIH-2 to promote GluA1 receptor tetramerization, we used an AMPAR construct lacking the ECDs (GluA1-ΔECD) (Gan et al 2016). GluA1-ΔECD efficiently forms tetramers, possibly more efficiently than full length GluA1 (**Figure 5D**) (Gan et al 2016), making it impossible for CNIH-2 co-expression to further enhance tetramerization. However, there is a slight, but notable upshift in molecular mass in the presence of CNIH-2 (**Figure 5D**), suggesting that the receptor can form a complex with the auxiliary subunit even without its extracellular domains. To assess whether CNIH-2 enhances the tetrameric stability of the truncated construct, we again took advantage of the lurcher mutation (A636T), which destabilized the tetramer formed by GluA1-ΔECD (**Figure 5D**). Co-expression of CNIH-2 significantly enhanced the tetrameric fraction, indicating that even without the ECD, CNIH-2 still enhances tetramerization (**Figure 5D**).

In summary, these results indicate that neither the CTD, nor the extracellular ATD and LBD of GluA1 is required for CNIH-2 to enhance receptor tetramerization. Taken together, our experiments suggest that interactions between the TMD of GluA1 and CNIH-2 can promote the process of AMPAR biogenesis. These experiments also suggest that the effect of CNIH-2 on receptor assembly is not dependent on mutations within the TMD, but rather reflects the stability of the tetrameric complex.

### Direct attachment of TARP γ-2 to GluA1 enhances receptor tetramerization

TARP γ-2 has no effect on AMPAR tetramerization (**Figures 1-4**). TARP γ-2 may not be able to stabilize the tetramer or does not interact with GluA1 in the early stages of AMPAR tetramerization and associates later during the AMPAR biogenesis process. We therefore took advantage of a construct, where the N-terminus of the auxiliary subunit is conjoined to the C-terminus of GluA1, referred to as the GluA1-TARP γ-2 tandem (Shi et al 2009, Miguez-Cabello et al 2020). The expression of wildtype GluA1-TARP γ-2 tandem predominantly and more efficiently forms tetramers in comparison to wildtype GluA1 alone (**Figure 6A**). To further verify this observation, we introduced the G816A mutation, which disrupts tetramerization (**Figure 2**), into the GluA1-TARP γ-2 tandem. As shown previously (**Figure 2C**), GluA1(G816A) co-expressed with TARP γ-2 alone had no effect on the tetramer fractions. In contrast, GluA1(G816A)-TARP γ-2 tandem showed almost complete tetramerization (**Figure 6B**).

**Figure 6.**
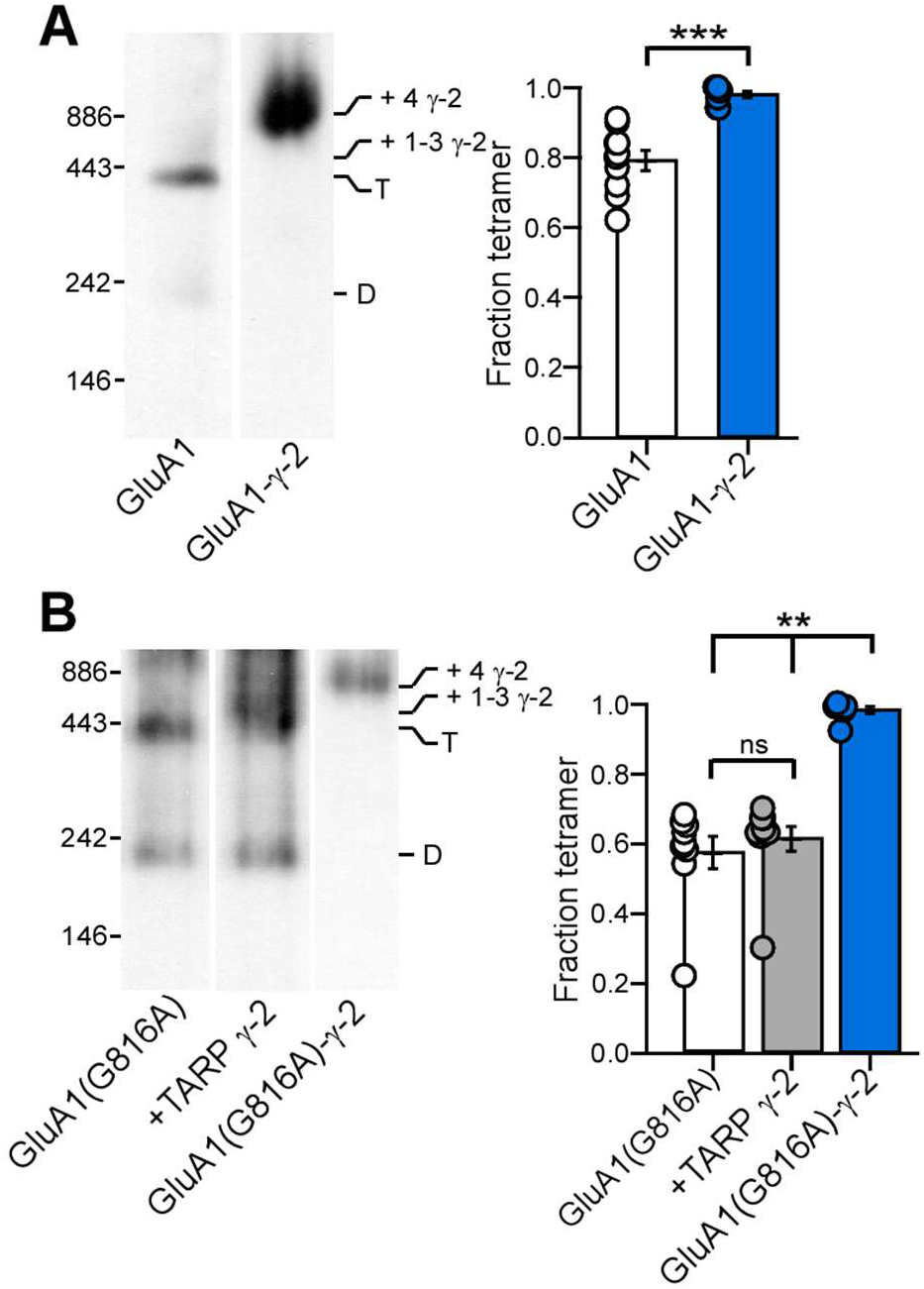
TARP γ-2 directly attached to GluA1 can enhance tetramerization. **(A)** BN-PAGE (*left panel*) and normalized tetramer fractions (*right panel*) of GluA1 or a construct where TARP γ-2 is directly attached to GluA1 (GluA1-γ-2). **(B)** BN-PAGE (*left panel*) and normalized tetramer fractions (*right panel*) for GluA1(G816A) expressed alone or with TARP γ-2 (+γ-2) or the GluA1-γ-2 tandem construct where the G816A mutation has been introduced (GluA1(G816A)-γ-2). Quantification of oligomeric states (left to right): n = 10, 8 (**A**); 9, 10, 7 (**B**). Number of gels (left to right) and significance (*t-test* or *one-way ANOVA with post-hoc Tukey test*) of tetramer fraction relative to control: 10, 8, *p < 0*.*001* (**A**); 9, 10, 7; G816A vs γ-2, *p = 0*.*71*; G816A vs tandem, *p = 0*.*001*; γ-2 vs tandem, *p = 0*.*001* (**B**). In plots, significance is indicated (*\*\*p < 0*.*01*, or *\*\*\*p < 0*.*001; ns*, not significant).

Additionally, the tetramer band of GluA1(G816A) co-expressed with TARP γ-2 appeared at a lower molecular mass than that of the GluA1(G816A)-TARP γ-2 tandem that is predicted to have a 4:4 stoichiometry (**Figure 6B**). This suggests that TARP γ-2 demonstrates a sub-maximal stoichiometry (4:1, 2, or 3) when co-expressed in HEK 293 cells at 1:1 ratio.

### CNIH-2 enhances surface GluA1 expression to a greater extent

To investigate whether the increased tetramerization of GluA1 leads to increased surface trafficking, and consequently higher current amplitudes, we performed whole-cell patch clamp experiments on HEK 293T cells expressing GluA1 with or without CNIH-2 or TARP γ-2. We were interested in the number of receptors on the membrane and not the effect of the auxiliary subunits on receptor kinetics, so we recorded currents in 15 μM cyclothiazide to minimize desensitization (Sun et al 2002, Milstein & Nicoll 2008). In preliminary experiments, wild type GluA1 showed very high current amplitudes making any increase in functional surface receptors with co-expression of auxiliary subunits difficult to identify. We therefore took advantage of GluA1(G816A), which shows reduced current amplitudes relative to wild type GluA1 in the absence of auxiliary subunits, but like GluA1, its tetramerization is enhanced by CNIH-2 but not by TARP γ-2 (**Figures 1, 2**).

GluA1(G816A) showed reduced current amplitudes that were strongly enhanced by CNIH-2 and TARP γ-2 (**Figure 7A**). In terms of current density, the amplitudes were equally increased for CNIH-2 and TARP γ-2 (**Figure 7B**). However, TARP γ-2 has a much stronger effect on enhancing GluA1 gating (*P*_*open*_) than CNIH-2 (Coombs et al 2012). We therefore derived a ‘Relative index of channels’ as a measure of density of GluA1 channels on the cell surface as outlined in Materials and Methods (**Figure 7C**), assuming that auxiliary subunits have the same effect on the channel properties of mutant and wild type GluA1 receptors. Based on this index, we found that CNIH-2 significantly enhanced surface expression of GluA1(G816A) relative to the control as well as relative to TARP γ-2. These results support the idea that enhanced receptor tetramerization leads to a functional effect on receptor membrane expression.

**Figure 7.**
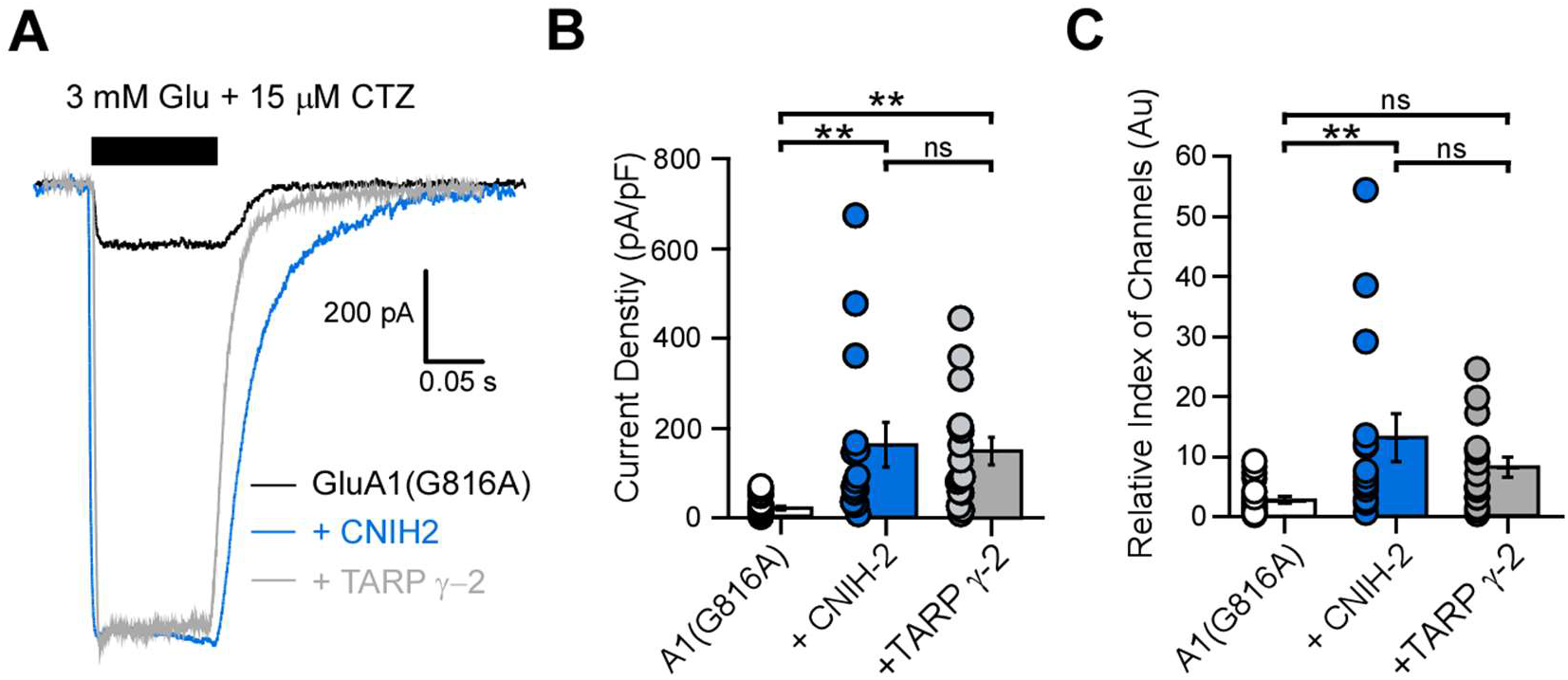
CNIH-2 enhances GluA1 membrane currents. **(A)** Whole-cell recordings of GluA1(G816A) expressed alone (black trace) or with auxiliary subunits (blue trace for CNIH-2; gray trace for TARP γ-2) in HEK293T cells. Solid bar indicates fast-agonist application of 3 mM glutamate. 15 μM CTZ was present throughout the recording. **(B)** Quantification (mean ± SEM) of steady state current amplitudes for GluA1(G816A) expressed alone or with auxiliary subunits (n = 22, 14, 16) normalized to membrane capacitance (C_m_, in pF) as an index of cell size. Significance (*one-way ANOVA with post-hoc Tukey test*): G816A vs CNIH-2, *p = 0*.*0048*; G816A vs γ-2, *p = 0*.*0074*; CNIH-2 vs γ-2, *p = 0*.*96*. **(C)** Quantification of relative index of channels of GluA1(G816A) expressed alone or with auxiliary subunits (n = 22, 14, 16). See Materials and Methods for normalization process. Significance (*one-way ANOVA with post-hoc Tukey test*): G816A vs CNIH-2, *p = 0*.*0055*; G816A vs γ-2, *p = 0*.*17*; CNIH-2 vs γ-2, *p = 0*.*34*. Significant differences of relative channel index are indicated (**p < 0.01; ns, not significant).

To further assess whether enhanced tetramerization would lead to enhanced surface expression, we transfected Neuro2A cells with GluA1(G816A) tagged with a C-terminal GFP either alone or with auxiliary subunits (**Figure 8A**). To assay surface expression, we measured integrated fluorescence of AMPARs present on the surface. Co-expression of CNIH-2 increased surface GluA1(G816A) receptors to a greater extent than TARP γ-2 expression (**Figure 8B**). On the other hand, the total protein level of GluA1(G816A) is not significantly altered by co-expression with either CNIH-2 or TARP γ-2 (**Figure 8C**). Together, these data indicate that CNIH-2 carries this distinct ability to increase AMPAR tetramers and enhances the forward trafficking and cell surface expression even when tetramer stability is attenuated.

**Figure 8.**
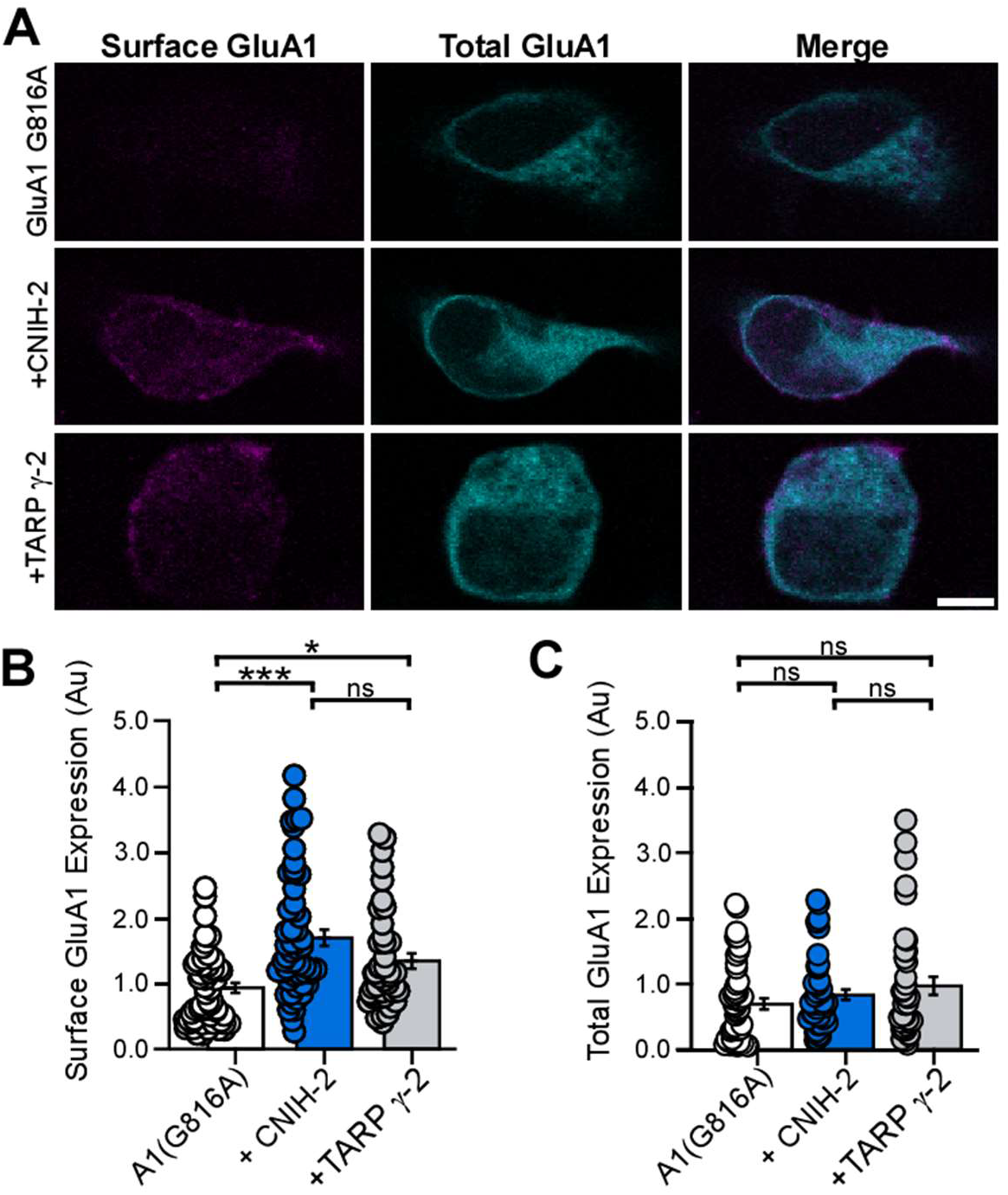
CNIH-2 enhances GluA1 surface expression. **(A)** Immunocytochemistry of Neuro2A cells co-transfected with GluA1(G816A)-eGFP alone or with auxiliary subunits, CNIH-2 or TARP γ-2. Surface GluA1 were immunostained using anti-GluA1 under non-permeabilized conditions. Scale bar represents 5 μM. **(D)** Quantification (mean ± SEM) of surface GluA1 expressed alone or with auxiliary subunits. Surface expression is normalized to GluA1(G816A) alone. GluA1(G816A), n = 52 cells; GluA1(G816A) + CNIH-2, n = 54 cells; GluA1(G816A) + TARP γ-2, n = 42 cells from 3 independent experiments. Significance (*one-way ANOVA with post-hoc Tukey test*): G816A vs CNIH-2, *p < 0*.*001*; G816A vs γ-2, *p = 0*.*028*; CNIH-2 vs γ-2, *p = 0*.*068*. **(E)** Quantification of total GluA1-eGFP expressed alone or with auxiliary subunits. Surface expression is normalized to GluA1(G816A) alone. GluA1(G816A), n = 48 cells; GluA1(G816A) + CNIH-2, n = 49 cells; GluA1(G816A) + TARP γ-2, n = 39 cells from 3 independent experiments. Significance (*one-way ANOVA with post-hoc Tukey test*): G816A vs CNIH-2, *p = 0*.*57*; G816A vs γ-2, *p = 0*.*14*; CNIH-2 vs γ-2, *p = 0*.*60*. Significant differences of are indicated (*p < 0.05, ***p < 0.001; or ns, not significant).

## Discussion

Assembly of AMPAR into a functional tetramer is required for the trafficking of AMPARs to the membrane surface (Gan et al 2015, Greger et al 2017, Jacobi & von Engelhardt 2020, Schwenk & Fakler 2020). In this study, we find that auxiliary subunits differentially influence AMPAR tetramerization. Specifically, cornichon homologs, CNIH-2 and CNIH-3, promote tetramerization of GluA1 AMPARs in a heterologous expression system (**Figure 1**), whereas for GluA2 this action is specific to CNIH-2 (**Figure 3**). While these effects were prominent in wild type GluA1 and GluA2, we also took advantage of a subtle mutation in the M4 transmembrane segment that attenuate tetramerization to further verify these outcomes (**Figures 2-4**). In contrast, other auxiliary subunits, including TARP γ-2, TARP γ-8, and GSGL-1, do not have any notable effect on receptor tetramerization. Hence, the differential expression of auxiliary subunits and their interaction with receptors in the ER may represent a distinct mechanism to control their availability for membrane expression.

CNIH-2 enhances tetramerization of both GluA1 and GluA2 homomers (**Figure 1-3**), suggesting that this effect is unlikely to be subunit-specific and applies to AMPARs in general. These results are consistent with the proposed role of CNIHs as ER chaperones based on their evolutionary origin as ER cargo exporters and their ER-Golgi localization in mammalian cells (Bokel et al 2006, Harmel et al 2012, Herring et al 2013). The loss of CNIH-2 leads to a loss of surface AMPARs (Harmel et al 2012, Herring et al 2013, Schwenk et al 2019), and CNIH-2 is well-known to promote ER exit of AMPARs (Shi et al 2010, Harmel et al 2012, Brockie et al 2013, Schwenk et al 2019). Our results further support CNIHs as a fundamental component of the AMPAR assembly process compared to other auxiliary subunits. Further studies are required to clarify how the role of CNIHs as an early promoter of AMPAR biogenesis impact synaptic function and plasticity.

The actions of CNIH-2 and CNIH-3 are not identical. Compared to CNIH-2, CNIH-3 enhances GluA1 tetramerization but appears less effective (**Figures 1 & 2**) and has no effect on GluA2 tetramerization (**Figure 3**). Knockout of CNIH-2 in excitatory neurons leads to a dramatic loss of surface GluA1-containing AMPARs, potentially consistent with CNIH-2 enhancing GluA1 tetramerization (Herring et al 2013, Liu et al 2018). Global knock-out (KO) of CNIH-3 in mice led to overall increases in GluA2-containing receptors in synaptosomes and no changes in GluA1 expression (Frye et al 2021). The phenotype of CNIH-3 KO could arise from a compensatory effect of CNIH-2 or the result of sex-related effects on CNIH-3 function (Frye et al 2021). Despite the difference in biogenesis, CNIH-2 and CNIH-3 have comparable effects on gating properties (Coombs et al 2012). The basis for the difference in receptor assembly between CNIH-2 and CNIH-3 is unknown.

It is unknown whether tetramerization is a limiting factor of AMPAR trafficking in neurons. It is possible that instead of increasing the overall size of the tetramer pool, CNIHs subtly shift the composition of the tetramers by selectively promoting the assembly of specific AMPAR subunit combinations. Indeed, the trafficking effect of CNIH-2 appears to be splice variant-dependent, and the loss of CNIH-2/3 from hippocampal neurons specifically impacts the surface expression of GluA1-containing receptors (Harmel et al 2012, Herring et al 2013). However, whether any of these effects are caused by a modulation of the tetramerization process is unclear.

CNIH-2 and CNIH-3 interact with AMPARs via the LBD, TMD and the LBD-TMD linkers of GluA2 (Shanks et al 2014, Hawken et al 2017, Nakagawa 2019). Using truncated constructs lacking entire structural domains, we show that neither the ECD (LBD & ATD) nor the intracellular CTD of GluA1 is required for CNIH-2 to enhance tetramerization (**Figure 5**). The interface at the transmembrane level of CNIHs and AMPARs are most likely responsible for the ability of CNIHs to promote AMPAR tetramers. Intriguingly, the interaction of the extracellular loop region of CNIHs and the LBD of AMPARs is required for functional modulation but does not seem important for enhanced receptor tetramerization (Shanks et al 2014). Hence, CNIHs may assist in receptor tetramerization by stabilizing certain inter-subunit interactions within the AMPAR TMD, such as those between the M4 segment and inner channel core (Salussolia et al 2013). Alternatively, CNIH-2 could exert its effect by interacting with and recruiting components of the AMPAR ER interactome such as FRRS1L, which in turn helps stabilize the tetramer formation in the ER (Rubio & Wenthold 1999, Brechet et al 2017).

Glutamate receptor tetramerization is a prerequisite for ER exit (Greger et al 2003, Nakagawa 2010, Salussolia et al 2013, Gan et al 2016, Schwenk et al 2019). The effect of CNIH-2 on initiation of ER exit is in part due to increased availability of fully assembled tetrameric complexes (Fleck 2006). Our experiments, however, suggest that in heterologous cells the abundance of tetramers in the ER is not a limiting factor for the number of surface-expressed receptors, since co-expression with TARP γ-2 was able to enhance the surface trafficking of GluA1(G816A) without rescuing the defect in tetramerization (**Figure 8B**). Both CNIH-2 and TARP γ-2 increase the surface expression of receptors in heterologous cells and primary neurons (Kato et al 2010a, Shi et al 2010, Harmel et al 2012). Our experiments in HEK293 cells show that CNIH-2 and TARP γ-2 increase the current amplitudes for exogenously expressed GluA1(G816A) to a similar degree but most likely perform these effects through different mechanisms (**Figure 7A-B**). The enhancement of current amplitude by TARP γ-2 can be largely accounted for by its ability to boost single-channel conductance and open probability (Coombs et al 2012, Shelley et al 2012, Soto et al 2014). The slight enhancement in surface trafficking, as evidenced by the upward trend in relative index of channels presented in Figure 7C (albeit statistically insignificant due to high variability) and by the increased surface fraction shown in Figure 8B, may also contribute to the enhancement of current amplitude by TARP γ-2. In contrast, since CNIH-2 does not enhance open probability as prominently as TARP γ-2, its effect on current amplitude most likely reflects its ability to promote assembly and surface trafficking of GluA1(G618A) (**Figure 7C & 8B**). We speculate that the main mechanism by which CNIH-2 promotes surface trafficking by efficiency to promote ER exit of fully assembled tetramers. Accordingly, stepwise expression of CNIH-2 with AMPARs compared to TARP γ-2 is more efficient in promoting assembled receptors (Schwenk et al 2019). The significance of this difference in the context of AMPAR biogenesis suggests that CNIH-2 differs from other auxiliary subunits in the stage at which it alters AMPAR maturation.

In contrast to CNIH-2, TARP γ-2 has little effect on AMPAR tetramerization (**Figure 1-3**), despite the association of TARP γ-2 with receptors in the ER (Bedoukian et al 2006). Instead, a maximal (4:4) stoichiometry, achieved by expressing GluA1 as a tandem with TARP γ-2, is required for the effect to manifest (**Figure 6**). Electrophysiological evidence suggests that AMPAR/TARP complexes display variable stoichiometry depending on neuronal types and the presence of other auxiliary subunits (Shi et al 2009, Gill et al 2011, Hastie et al 2013, Miguez-Cabello et al 2020), a notion that is also supported by recent structural data of native receptors (Zhao et al 2019). One possible explanation is that the maximal stoichiometry of 4:4 between GluA1 and TARP γ-2 has a lower incidence and is dependent of expression levels of TARP γ-2 (Kim et al 2010, Hastie et al 2013, Miguez-Cabello et al 2020). It is possible that CNIHs are more likely to associate with AMPARs at a 4:4 stoichiometry than TARPs in HEK293 cells. Alternatively, the effect of CNIHs on AMPAR tetramerization, unlike that of TARPs, is independent of stoichiometry. Due to the inherent ambiguity of molecular mass measurement in native gels, and the unavailability of a viable GluA1-CNIH-2 tandem constructs, we could not determine whether CNIH-2 co-expressed with GluA1 in our experiments reached a maximal stoichiometry (Schagger et al 1994).

An alternative explanation of the differential effects between TARP γ-2 co-expression and γ-2 tandem constructs relates to the stage at which TARP γ-2 associates with AMPARs: when co-expressed in HEK293 cells, TARP γ-2 and AMPARs may only interact at later stages of biogenesis (e.g., in golgi complex). In contrast, when expressed as a tandem construct, TARP γ-2 and AMPARs are forced to interact in the ER, where AMPAR tetramerization occurs. We speculate that CNIH-2 was able to influence the tetramerization of GluA1(G816A) because it associates with the receptor at an earlier stage during receptor biogenesis compared to TARP γ-2 (Harmel et al 2012).

Auxiliary subunits play a key role in regulating AMPAR function at the surface, including channel gating and synaptic targeting (Bissen et al 2019). However, prior studies have not investigated how auxiliary subunits may regulate receptor assembly, before a receptor can reach the surface. Our study found that CNIHs can differentially regulate AMPAR assembly by increasing the fraction of tetramers. The early stages of AMPAR biogenesis can be disrupted in pathological conditions (Brechet et al 2017, Groc & Choquet 2020) and may be crucial to our understanding of certain neurological disorders. Overall, the results obtained in the study suggest that normal receptor assembly is regulated by auxiliary subunits and disruption to this role can lead to synaptic dysfunction suggested by the genetic risk of psychiatry and addiction susceptibility in CNIH-2 and CNIH-3 (Nelson et al 2016, Han et al 2018).

## Acknowledgments

This work was supported by a NIH RO1 grant from NINDS (NS088479) (LPW) including a NINDS Diversity supplement (NC), a NINDS F99 NS124126 (NC), and a Dr. W. Burghardt Turner Dissertation Fellowship (NC). We thank Drs. Stella Tsirka, Joshua Plotkin, Louis Manganas, Susumu Tomita, David McKinnon, Maurice Kernan, and Hiro Furukawa for helpful discussions and/or comments on the manuscript. We thank Drs. Bernd Fakler, Stephan Herlitze, and Susumu Tomita for providing cDNAs for various auxiliary subunits. We also thank Drs. Roger Nicoll and Yun Shi for providing the AMPAR-TARP γ-2 tandem constructs.

